## Supplemental Figures for "UV irradiation alters TFAM binding specificity and compaction of DNA"

**This file includes:**

Figures S1 to S19

Tables S1 to S2

SI References

**Table S1: Total number of TFAM complexes analyzed in Figure 4D.**

| **Category** | **15nM TFAM -UV** | **15nM TFAM +UV** | **30nM TFAM**  **-UV** | **30nM TFAM**  **+UV** |
| --- | --- | --- | --- | --- |
| Free DNA | 74 | 14 | 4 | 5 |
| Dispersed | 21 | 9 | 29 | 2 |
| Intermediate (tracts) | 36 | 20 | 47 | 19 |
| Punctate | 5 | 23 | 11 | 39 |
| **Total Number of DNAs** | N = 136 | N = 66 | N = 91 | N = 65 |

**Table S2: Sequences used for TFAM K_D_ measurements.**

| **Oligonucleotide name** | **Sequence (5’ – 3’)** |
| --- | --- |
| ND1_288 | CTC CAC ACT AGC AGA GAC CAA CCG AAC CCC CTT |
| COX2_229 | GCC CCC ATT CGT ATA ATA ATT ACA TCA CAA GAC |
| TRNT_10 | CTT GTA GTA TAA ACT AAT ACA CCA GTC TTG TAA |
| ND2_401 | CAA ATG GGC CAT TAT CGA AGA ATT CAC AAA AAA |
| ND3_92 | TTA GTA GCT ATT ACC TTC TTA TTA TTT GAT CTA |
| ND1_450 | ACT CAC CCT AGC ATT ACT TAT ATG ATA TGT CTC |
| ND6_87 | TTC CTA CAC TAT TAA AGT TTA CCA CAA CCA CCA |
| RNR2_619 | ATT GAT CCA ATA ACT TGA CCA ACG GAA CAA GTT |
| COX1_27 | AAG ACA TTG GAA CAC TAT ACC TAT TAT TCG GCG |
| ND4_473 | CGG CGC AGT CAT TCT CAT AAT CGC CCA CGG GCT |
| ND1_353 40mer | CAC AAA CAT TAT TAT AAT AAA CAC CCT CAC CAC TAC AAT C |
| ND1_353 | ATT ATT ATA ATA AAC ACC CTC ACC ACT ACA ATC |

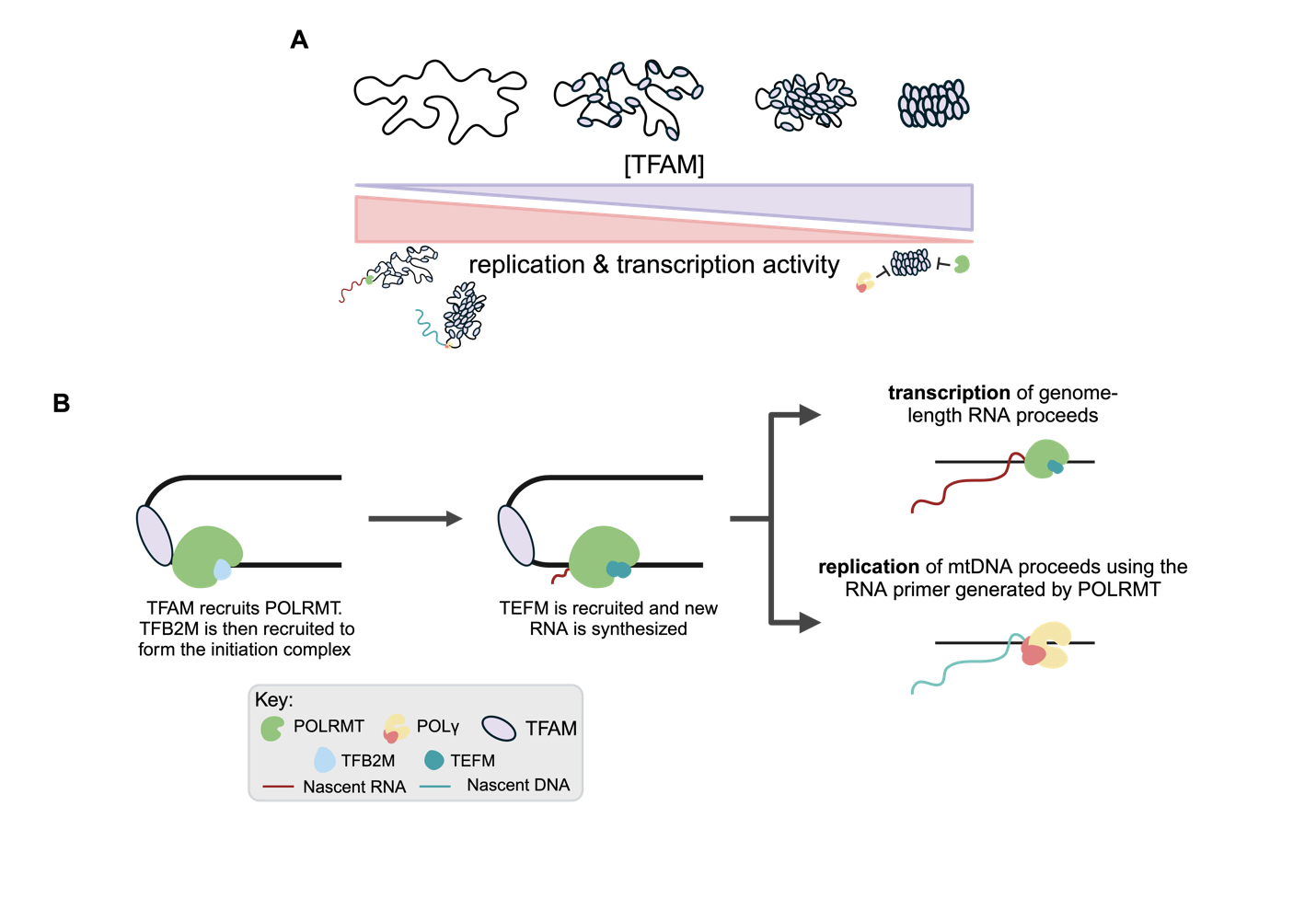
**Figure S1: TFAM roles in regulating the mitochondrial genome**. **A)** TFAM is one of the dominant proteins of the mitochondrial nucleoid and has been shown *in vitro* to be sufficient to coat and compact the genome. Compactional status of the mitochondrial nucleoid is associated with the activity of the nucleoid wherein loose and open genomes are accessible to polymerases and protein complexes required for transcription and replication, while compacted and closed off genomes are inaccessible and inactive. **B)** In its role as a transcription factor, TFAM serves to initiate both transcription and replication. TFAM binds upstream one of the transcription start sites and recruits mitochondrial RNA polymerase (POLRMT). Transcription factor B2 mitochondrial (TFB2M) is then recruited to form the initiation complex. Synthesis of RNA by POLRMT then proceeds with transcription elongation factor mitochondrial (TEFM). Transcription of genome-length RNA transcripts can then occur. Production of a short RNA primer by POLRMT allows for the replication of the mitochondrial genome by the replicative polymerase γ, comprising the catalytic subunit (POLG) and its accessory subunit (POLG2). Figure made in BioRender.

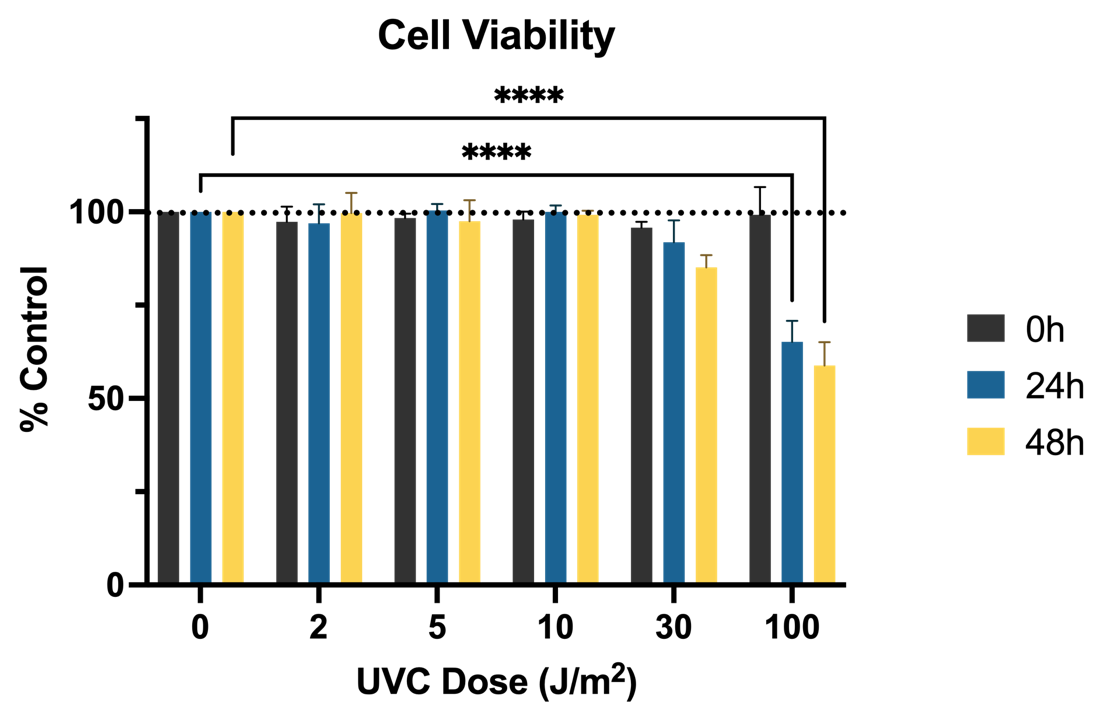

**Figure S2: Cell viability at 0-, 24-, and 48-hours following exposure to UVC**. Cell viability following exposure to UVC, normalized to the control for each replicate (n=3), determined using a resazurin cell viability assay. X-axis represents the UVC dose delivered to the cells in J/m^2^. Y-axis represents the fluorescent readout normalized as percent control. Data was analyzed via two-way ANOVA with a Dunnett’s correction for multiple comparisons to the control for each time point (dose: p=0.004, time: p<0.001, interaction: p=0.002, 24h 100 J/m^2^ p<0.0001, 48h 100 J/m^2^ p<0.0001).

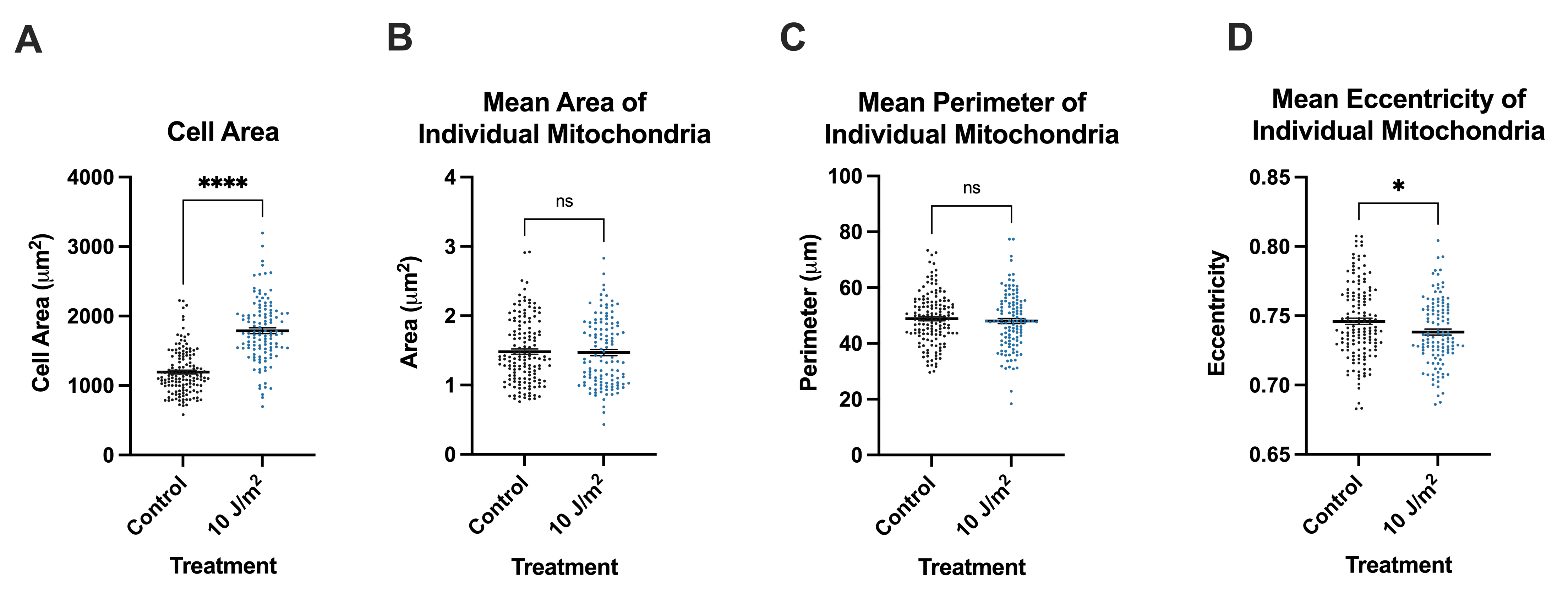

**Figure S3: Cell area and mitochondrial morphology analysis from live-cell imaging experiments.** For all panels, x-axes represent the treatment and data was analyzed via two-tailed unpaired t-test. Data includes at least n=30 cells per treatment group per imaging session and three distinct imaging sessions. **A)** Quantification of the total area per cell (p<0.0001). **B)** Quantification of the mean area of individual mitochondria within each cell (p=0.84). **C)** Quantification of the mean perimeter of individual mitochondria within each cell (p=0.45). **D)** Quantification of the mean eccentricity values of individual mitochondria within each cell (p=0.02).

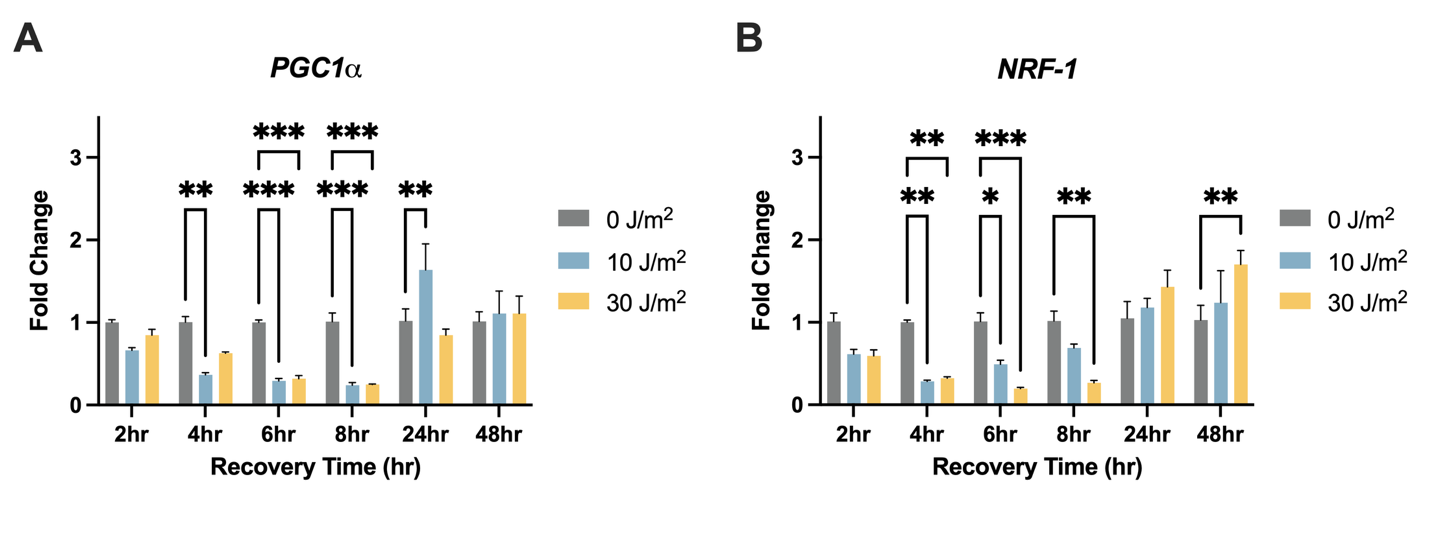

**Figure S4.** **UVC exposures alter gene expression of *PGC1α* and *NRF-1*.** For all both panels, x-axes represent recovery time, i.e., time following the UVC exposure, and y-axes represent fold change normalized to the control (0 J/m^2^) at each timepoint. Doses of UVC used were 0, 10, and 30 J/m^2^. All data was analyzed via two-way ANOVA with Dunnett’s post-hoc test for multiple comparisons. **A)** *PGC1α* expression level assessed via qPCR following UVC exposure (dose: p<0.0001, time: p<0.0001, interaction: p<0.0001). **B)** *NRF-1* expression level assessed via qPCR following UVC exposure (dose: p<0.0001, time: p=0.0022, interaction: p<0.0001).

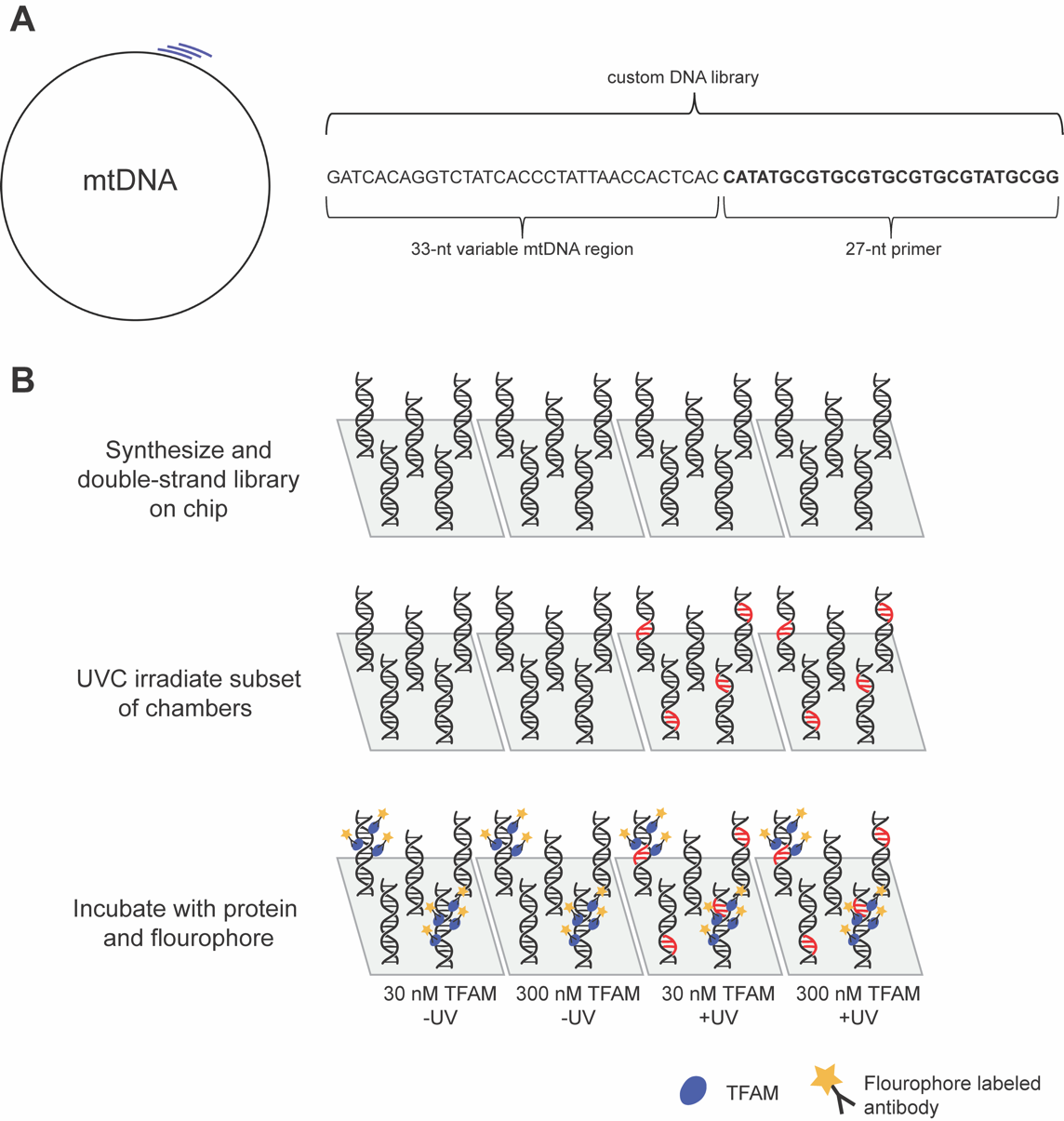

**Figure S5: Custom DNA library with full coverage of human mtDNA genome with a sliding window width of 2 nucleotides**. **A)** Schematic depicting the library generation. Each 60nt sequence contains a 33nt variable region from the human mitochondrial genome and a 27nt primer. To ensure full coverage of the mitochondrial genome, the variable region of the sequence was generated using a sliding window with a width of 2nt. **B)** The custom DNA library was synthesized and double-stranded on a chip. Two of the chambers were subjected to UVC irradiation to induce UVC associated lesions. Chambers were incubated with either 30 or 300 nM TFAM and a fluorophore labeled antibody. The fluorescent signal associated with bound protein for each DNA spot was determined using a microarray scanner.

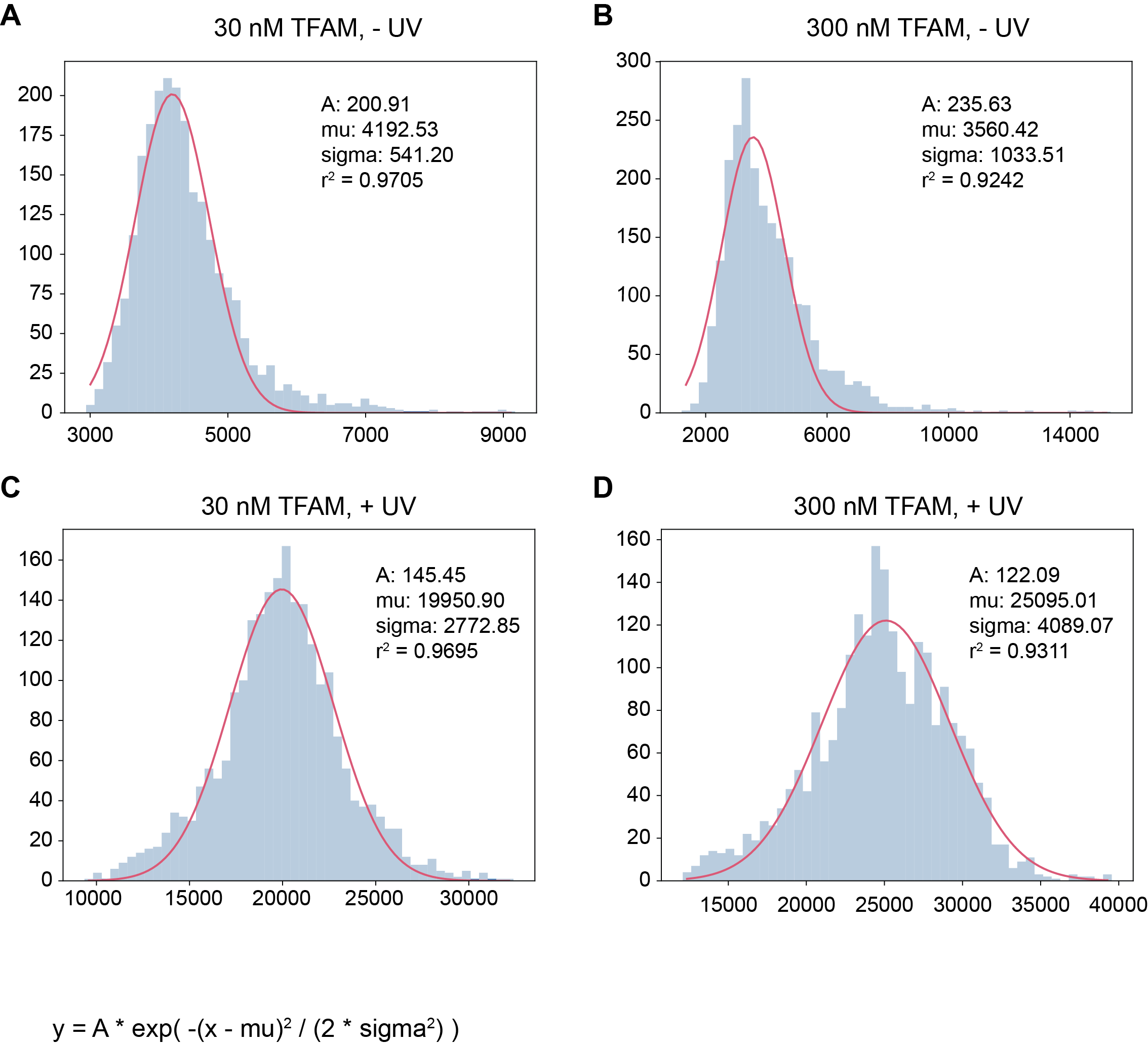

**Figure S6: Fluorescence intensity distribution of the bottom 116 non-mitochondrial sequences from a universal DNA-binding array used to calculate z-scores for each chamber.** X-axes represent the normalized median fluorescence intensity values and y-axes represent the count of sequences within each bin for 30 nM TFAM treatment without UVC irradiation (**A**), 300 nM TFAM treatment without UVC irradiation (**B**), 30 nM TFAM treatment with UVC irradiation (**C**), and 300 nM TFAM treatment with UVC irradiation (**D**). The red line is the Gaussian fit using the parameters in each plot and the equation below, where mu is the mean and sigma is the standard deviation.

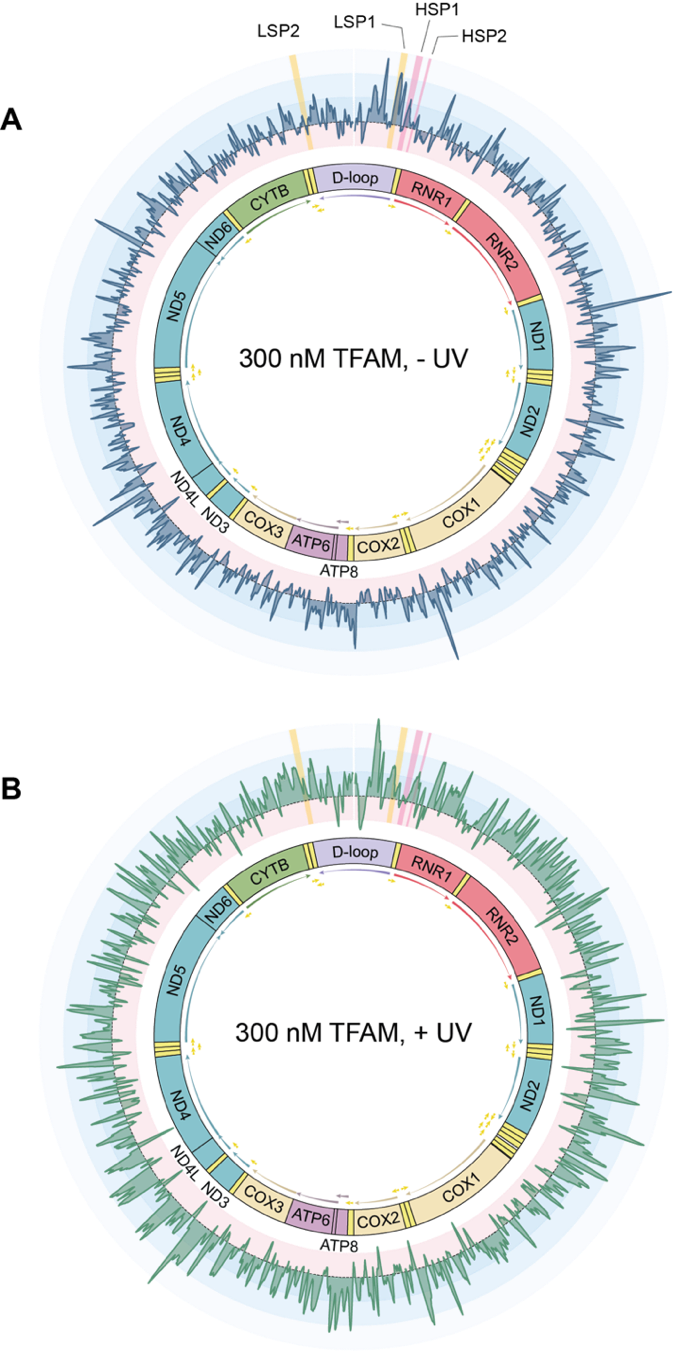

**Figure S7: Experiments performed at 300 nM TFAM concentrations demonstrate that TFAM has specific binding across the mitochondrial genome and exhibits a reduction in specificity in the context of UVC-irradiated DNA.** The median z-score plotted to the coordinate of the middle nucleotide of the variable mitochondrial region of the sequence for the non-UVC-irradiated chamber containing 300 nM TFAM (**A**) and the UVC-irradiated chamber containing 300 nM TFAM (**B**). The gene map of the mitochondrial genome is shown in the center. Z-score variation is color coded such that positive z-scores associated with high binding are in in blue, and progressively get lighter as the z-scores get higher. Negative z-scores associated with low binding are in red. Regions highlighted in yellow are the promoter sequences of the mitochondrial genome on the light strand (LSP1 and LSP2), while regions highlighted in pink are the promoter sequences on the heavy strand (HSP1 and HSP2)

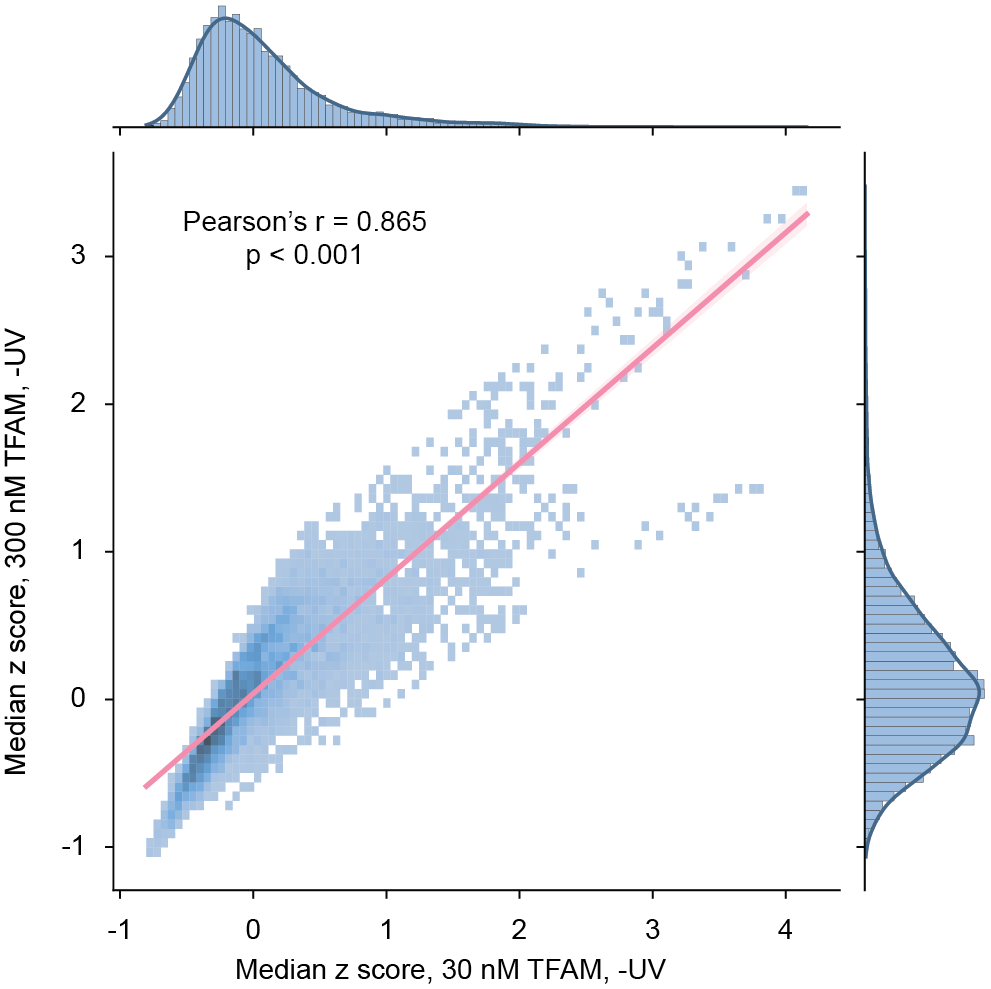

**Figure S8: Correlation between z-scores obtained at 30 nM and 300 nM TFAM.** The x-axis represents the median z-score for each probe in experiments performed at 30 nM TFAM and the y-axis represents the median z-score for each in experiments performed at 300 nM TFAM. Distributions to the right and above the graphs represent the distribution of z-scores for each concentration. The red line indicates the line of best fit. Data was analyzed via Pearson’s correlation.

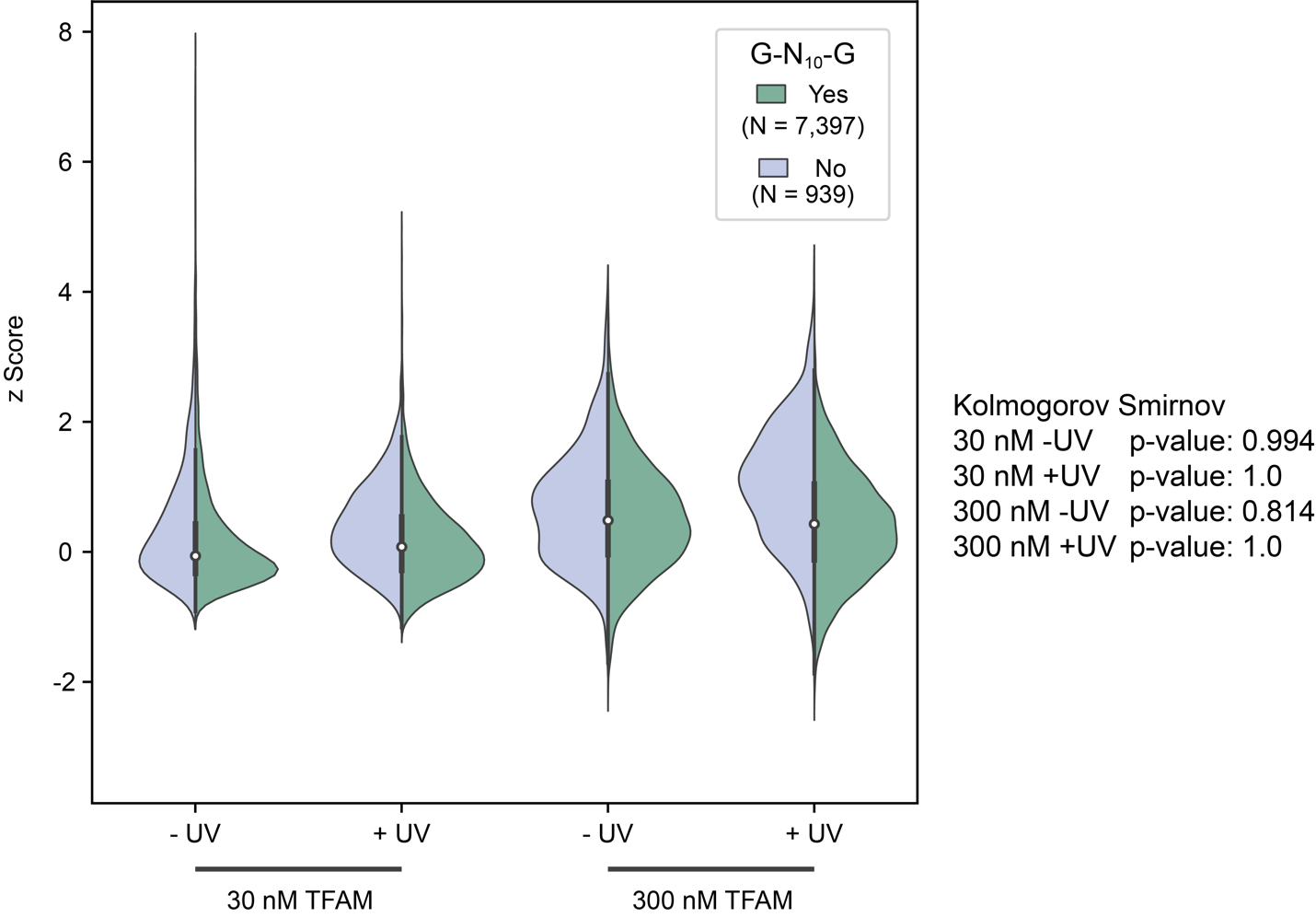

**Figure S9: Across all experiments, GN_10_G motifs are not enriched in high TFAM occupancy groups.** The y-axis represents the z-score for each sequence and the x-axis depicts the concentration of TFAM used in each experiment as well as the presence or absence of UVC exposure. The violin plots are color coded to represent whether a GN_10_G motif was present (green, n=7,397) or absent (blue, n=939) within each sequence. For all mitochondrial sequences on the array, sequences were classified as having a GN_10_G motif if the motif was present in the variable region and the data analyzed was the maximum z-score between the two orientations. Data was analyzed using a one-tailed Kolmogorov-Smirnov test where the null hypothesis was sequences containing GN_10_G motif have higher z-scores than those without.

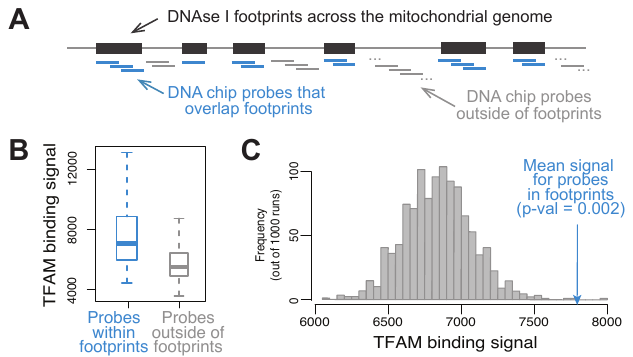

**Figure S10:** **DNAse I footprints across the mitochondrial genome are enriched for sites with high *in vitro* binding signals for TFAM**. **A)** For all DNase Genomics Footprinting (DGF) sites reported by Blumberg *et al*. (1), that were identified in >90% of analyzed human samples, we selected all 33-bp mtDNA probes in our on-chip DNA library that were contained entirely within DGFs. For DGFs shorter than 33-bp, we selected the DNA chip probes that contained the entire DGF. **B)** The TFAM DNA-binding signal (*i.e.*, the fluorescence intensity measured on the DNA chip) was significantly higher at probes that overlap footprints (blue) vs. sites outside of footprints (grey); Mann-Whitney test p-value < 2.2x10^-16^. **C)** We also performed a randomization test where we shuffled the positions of the DGF sites 1,000 times (keeping the number of DGFs and the distribution of their lengths constant). For each randomization we repeated the analysis and found that in only 2 of 1000 random sample the mean TFAM binding was at least at large as in the real DGF data (p=0.002).

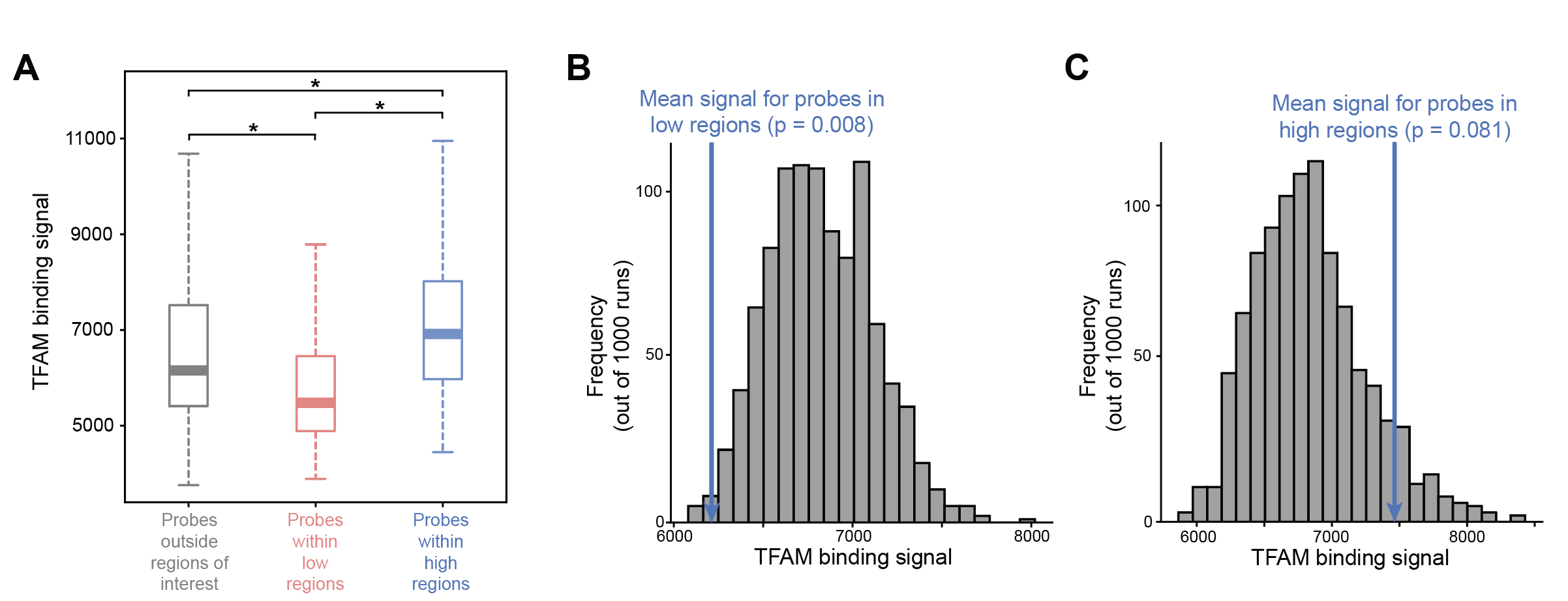

**Figure S11: Comparison between fiber-seq high and low TFAM binding regions and our high-density TFAM-DNA binding array data.** For high and low TFAM binding regions reported by Isaac *et al.* (2), determined using fiber-seq on linear mtDNA, we selected all 33-bp mtDNA probes in our on-chip DNA library that were contained entirely within the region. For regions shorter than 33-bp, we selected the DNA chip probes that contained the entire region. **A)** For probes within the low binding regions, we observe lower TFAM DNA-binding signal (*i.e.*, the fluorescence intensity measured on the DNA chip) (red) and for probes within the high binding regions, we observe moderate to high levels of TFAM DNA-binding signal (blue) than all other probes that were not within the high and low TFAM regions reported by Isaac *et al.* Data was analyzed via Mann-Whitney test (all other probes vs. low regions p-value < 3.0x10^-18^; all other probes vs. high regions p-value < 8.9x10^-05^; low regions vs. high regions p-value < 2.8x10^-13^). We also performed a randomization test where we shuffled the positions of both the low **(B)** and high **(C)** TFAM binding regions 1,000 times (keeping the number of regions and the distribution of their lengths constant). For the low binding regions, only 8 of the 1000 random samples had binding signal less than the mean for the probes within the low regions (p=0.008). However, for the high binding regions, 81 of the 1000 random samples had binding signal higher than the mean for the probes within the high regions (p=0.081).

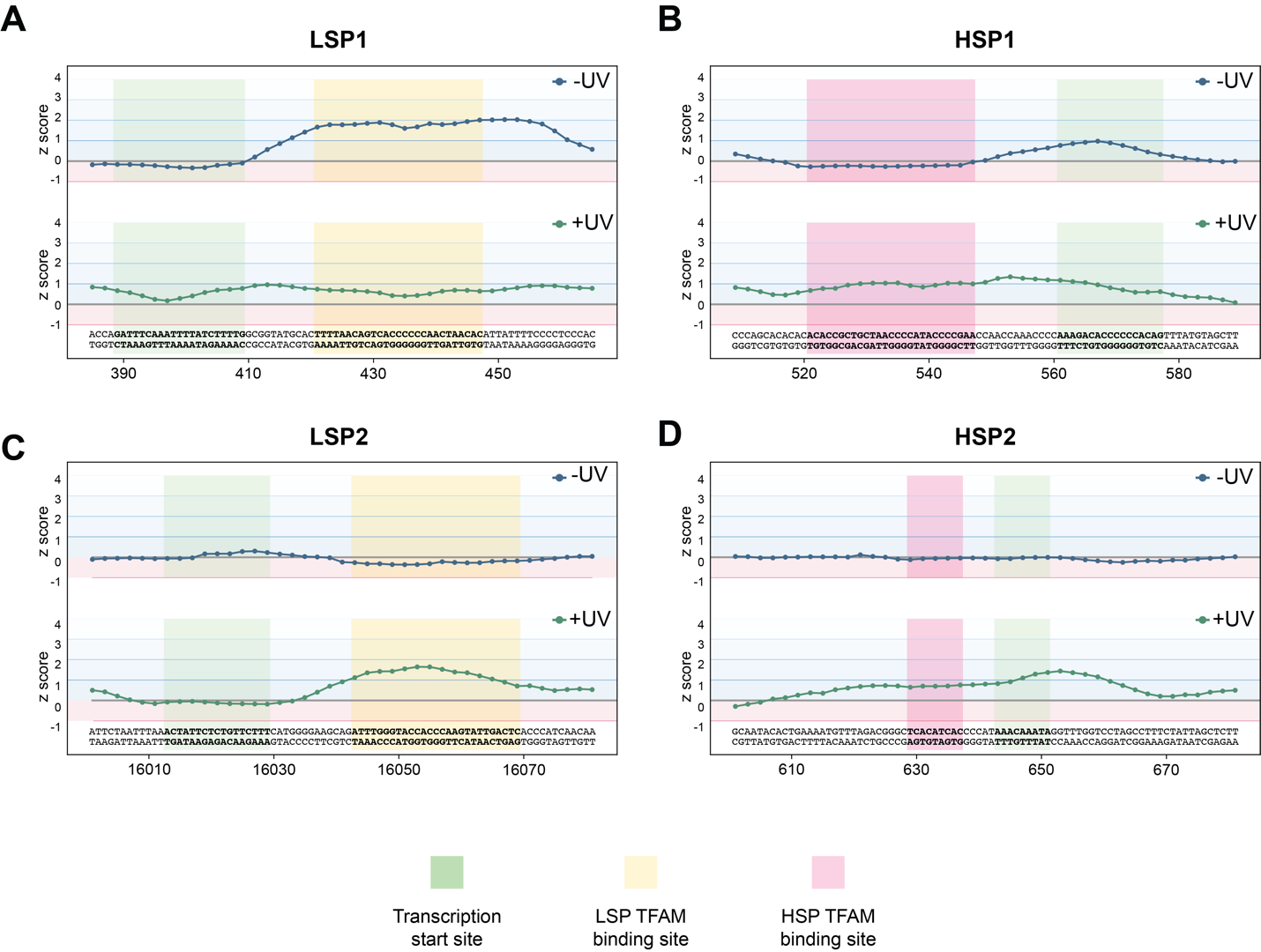

**Figure S12:** **High resolution view of TFAM binding to promoter sequences and shift in z-scores following UVC-irradiation.** For panels A-D, the median z-score is plotted to the coordinate of the middle nucleotide of the variable mitochondrial region of the sequence for the non-UVC-irradiated chamber containing 30 nM TFAM (**top panels, in blue**) and the UVC-irradiated chamber containing 30 nM TFAM (**bottom panels, in green**). Z-score variation is color coded such that positive z-scores associated with high binding are in in blue and negative z-scores associated with low binding are in red. Panels represent the shaded regions in Figure 3 for LSP1 **(A)**, HSP1 **(B)**, LSP2 **(C)**, and HSP2 **(D)**. Regions highlighted in yellow are LSP TFAM binding sites, regions highlighted pink are HSP TFAM binding sites, and regions highlights in green are transcription start sites.

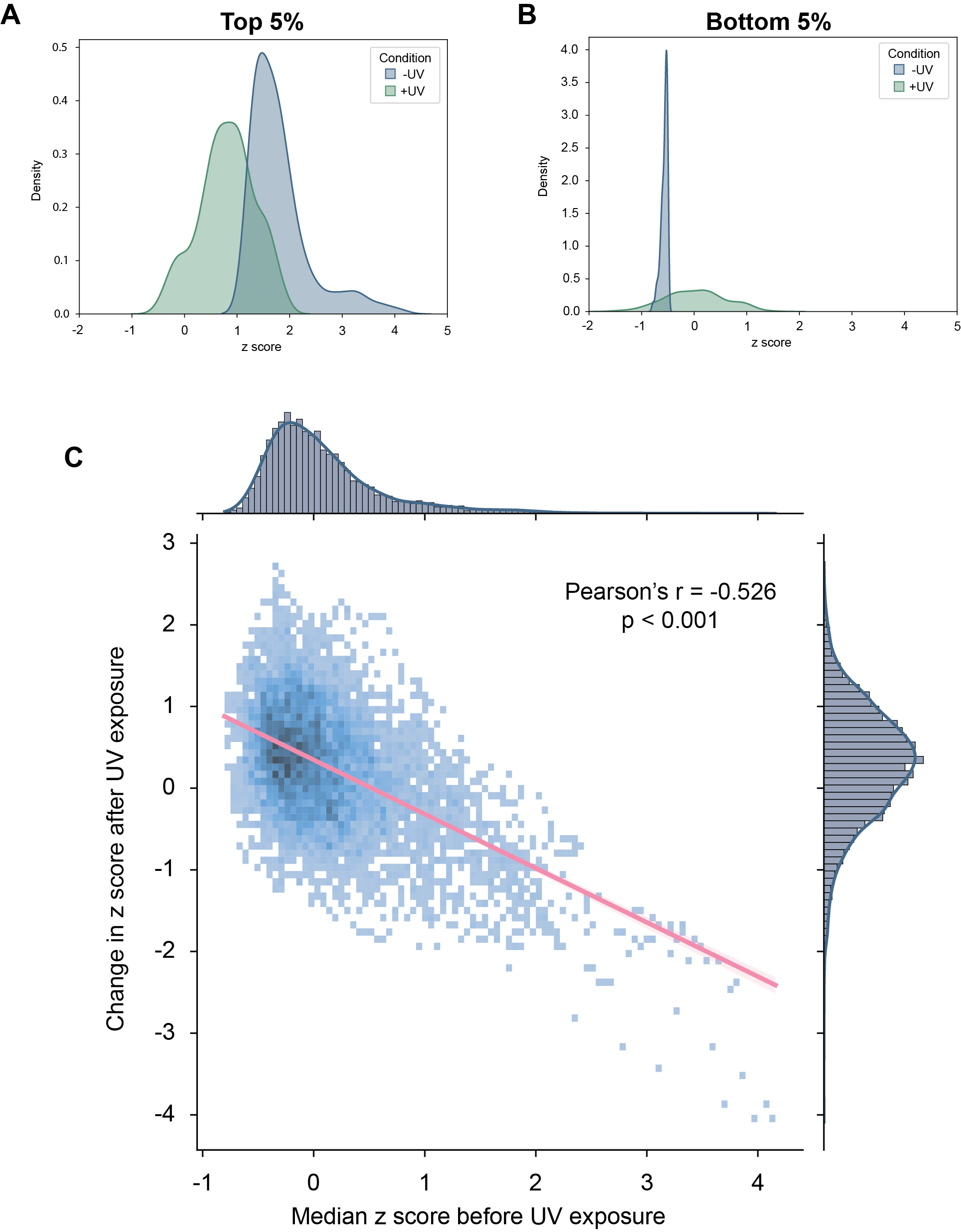

**Figure S13: UVC-irradiation is associated with a reduction in TFAM sequence specificity as weakest binders become tighter and tightest binders become weaker.** Panels A and B are kernel density estimates of the probes in the top 5% of z-scores **(A)** and the bottom 5% of z-scores **(B)** in the context of non-damaged DNA and UVC-irradiated DNA. X-axes represent the median z-score of the probe and y-axes represent the probability density. Note differences in y-axes between panels A and B. **(C)** The X-axis represents the median z-score for each probe in experiments performed at 30 nM TFAM and the y-axis represents the change in z-score for each probe between non-UVC and UVC-irradiated experiments. The distribution plot above of the graph indicates the distribution of median z-score values in the non-damaged probes performed at 30 nM TFAM. The distribution plot to the right of the graph indicates the distribution of the difference in z-scores calculated between non-UVC and UVC-irradiated experiments. The red line indicates the line of best fit. Data was analyzed via Pearson’s correlation.

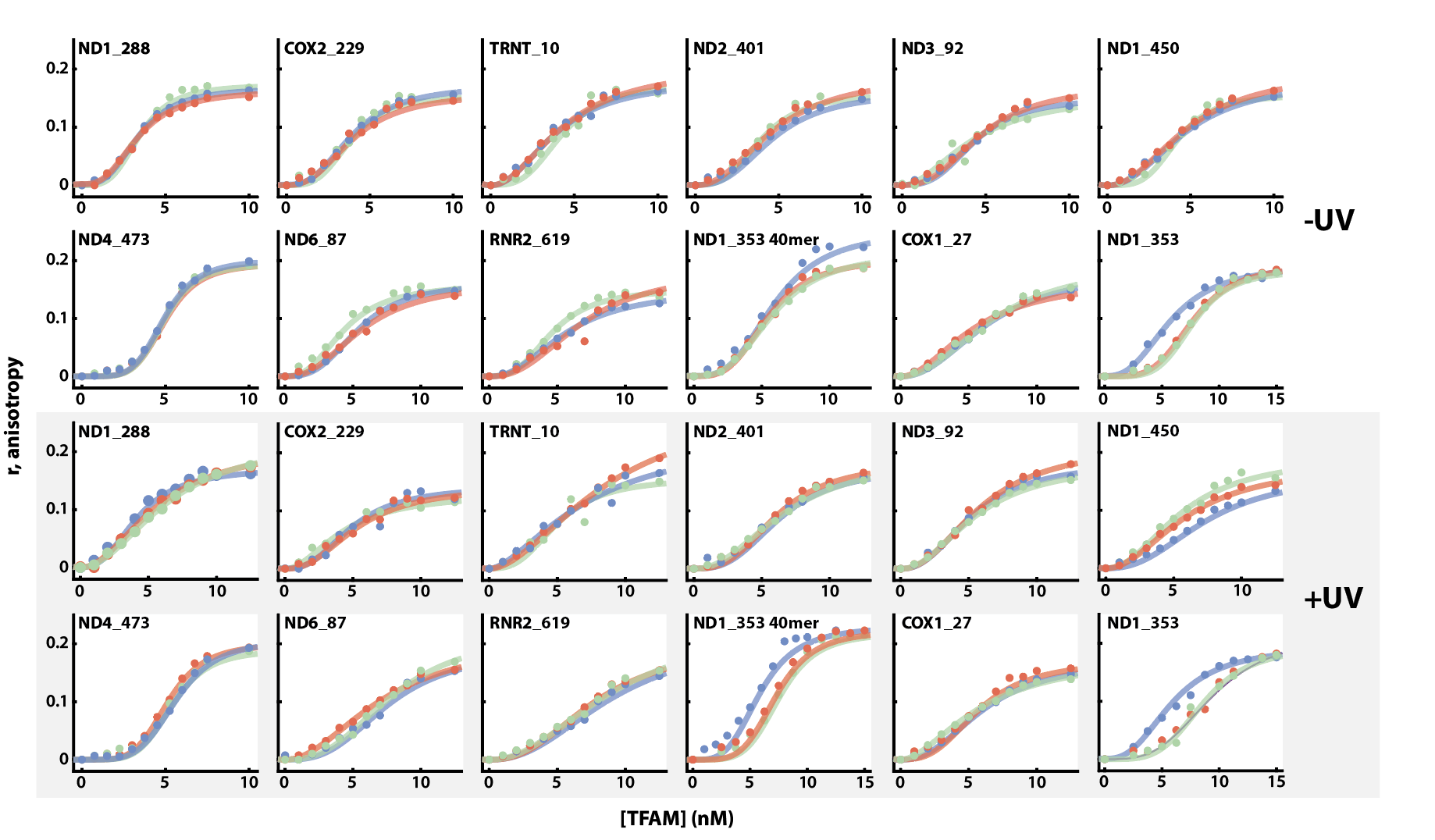

**Figure S14: Individual anisotropy plots for all sequences tested.** For all graphs, the x-axes represent the TFAM concentration (nM) and the y-axes represent anisotropy. The three lines on each graph represent three replicates performed. Graphs in the top half (white) are the sequences tested without UV exposure. Graphs in the bottom half (grey) represent sequences that were first irradiated with UVC. Sequences can be found in Table S2. K_D_ values and n values can be found in Table 1.

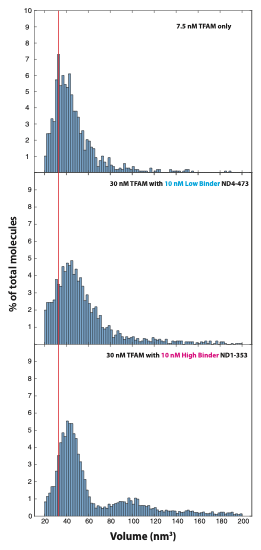

**C**

**B**

**A**

**Figure S15: Histogram plots of volumes for TFAM with and without DNA oligonucleotides. A)** Top panel is a histogram plot for the volumes of TFAM protein only at 7.5 nM. The red line is aligned with the peak at 33 nm^3^ in the TFAM only histogram to help show the shift in peaks for the TFAM with DNA oligonucleotides in the two bottom panels. **B)** Middle panel is the volume distribution of a 30 nM TFAM incubated with a final concentration of 10 nM of the low binder oligonucleotide ND4-473. **C)** Bottom panel is the volume distribution of 30 nM TFAM in the presence of 10 nM high binding oligonucleotide ND1-353.

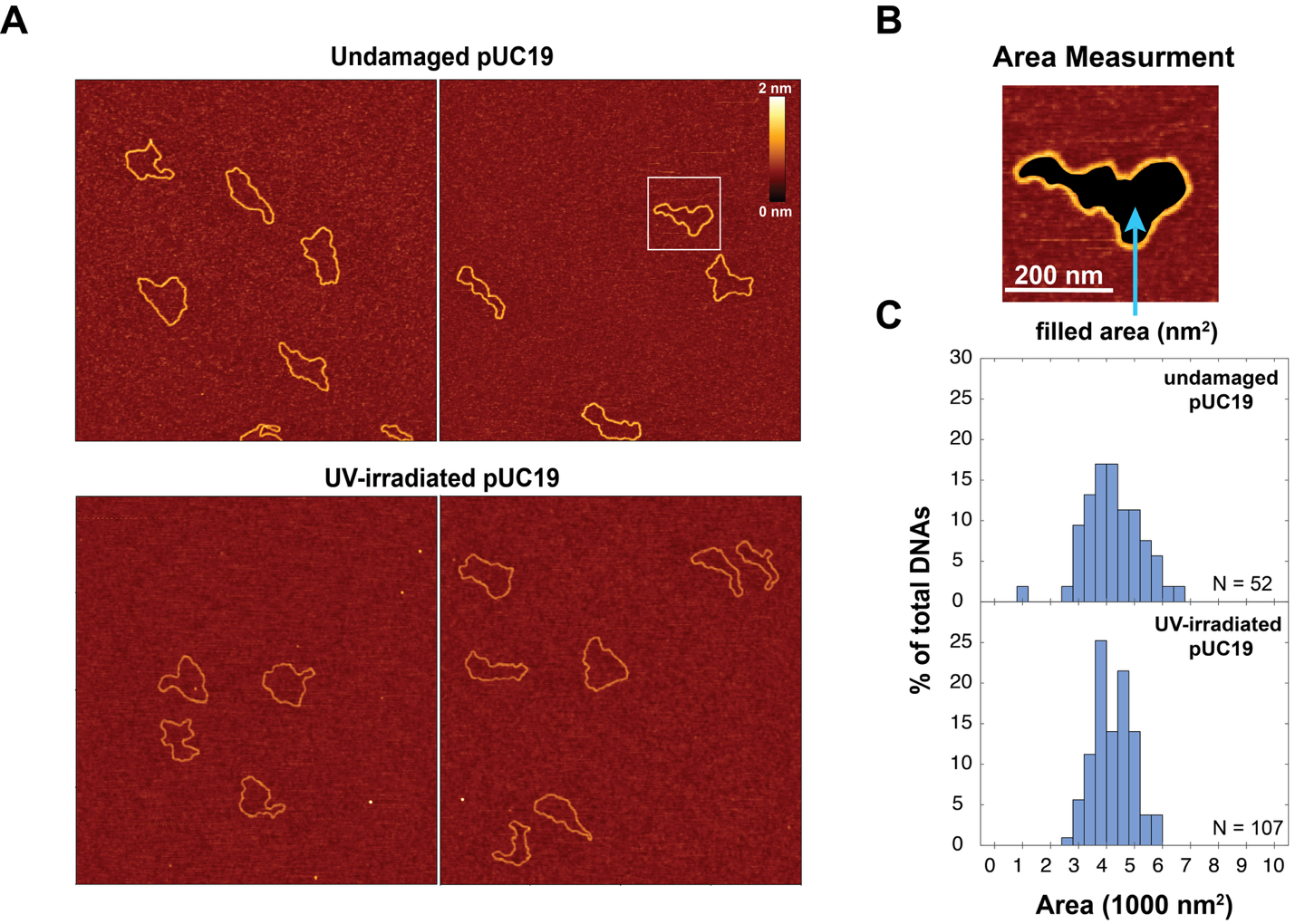
**Figure S16: Area quantification of undamaged and UV-irradiated pUC19 plasmids using AFM. A)** Representative AFM images of undamaged pUC19 (top) and UV-irradiated pUC19 (bottom). **B)** Overview of how area (nm^2^) of each plasmid was analyzed. The area is defined as the region encapsulated within the DNAs colored in black. **C)** Histogram plots of the areas for both undamaged pUC19 (top) and UV-irradiated pUC19 (bottom). The average area for the undamaged pUC19 is ~42,170 nm^2^ and the average area for the UV-irradiated pUC19 is ~42,200 nm^2^. Each AFM image is 2x2 μm and 512 x 512 pixels.

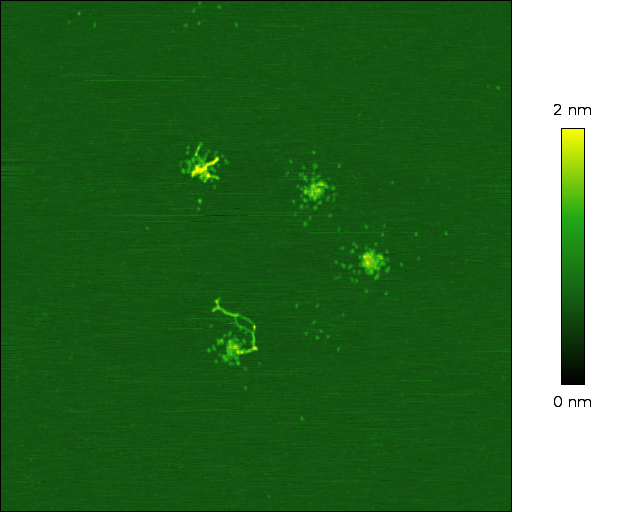

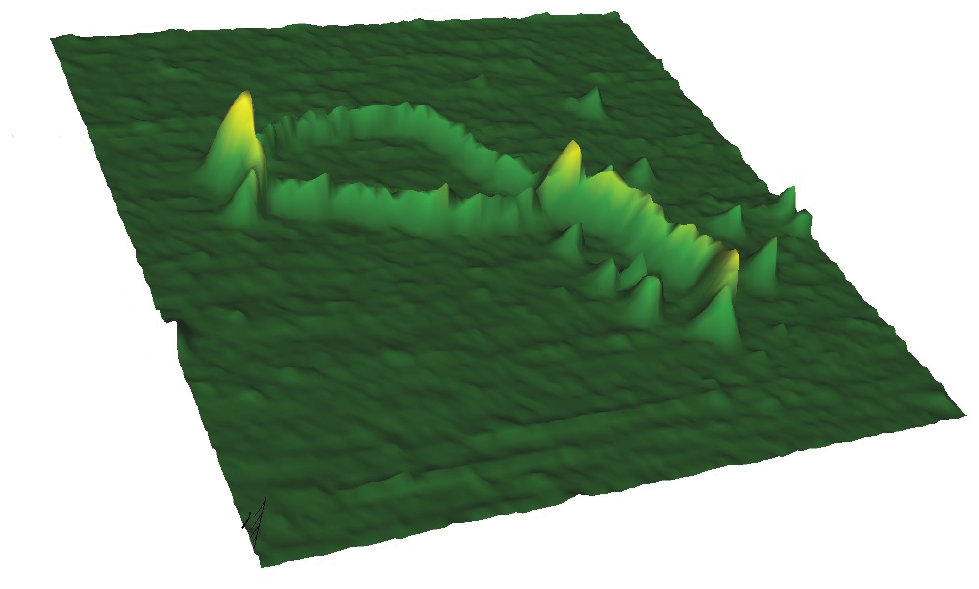

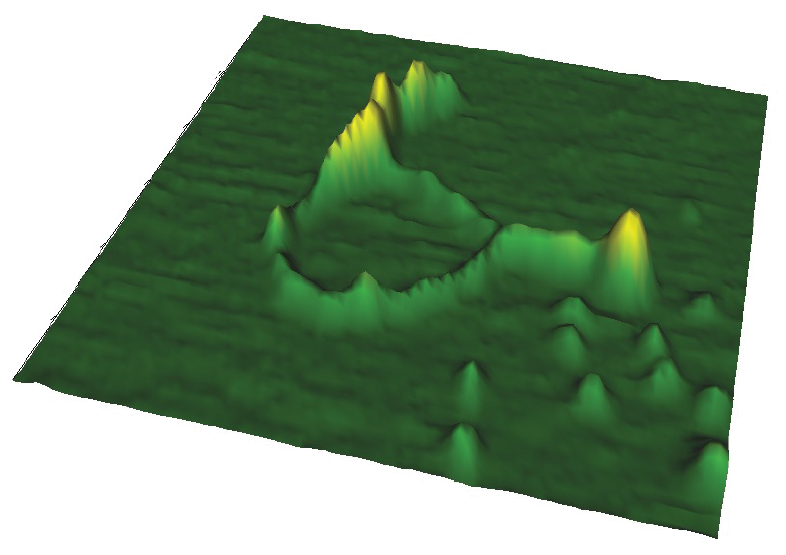
**Figure S17: 2D and 3D AFM images of TFAM tracts along DNA. A)** 2D AFM images with TFAM-DNA complexes, with white arrows pointing at the tracts of TFAM along the DNA (white scale bars are all 100 nm). **B)** 3D AFM images of two images in panel A (images labeled 1 and 2) showing the difference in areas along the DNA without protein and regions with DNA and protein (TFAM tracts). In the 3D AFM images it is clear to see that the areas with DNA only (no protein) are much lower in height (nm) whereas the areas with protein along the DNA (TFAM tracts) are higher in height.

no protein along DNA

2

1

2

1

**B**

tracts

tracts

no protein along DNA

tracts

no protein along DNA

**A**

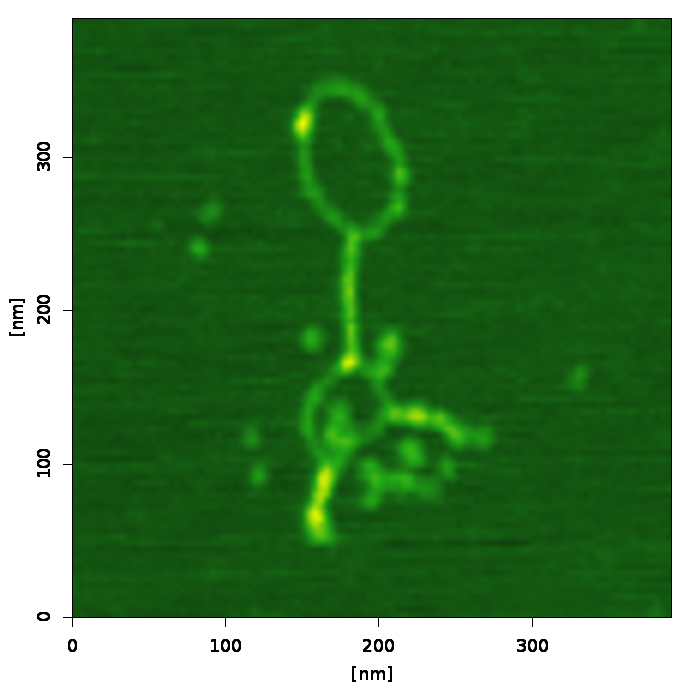

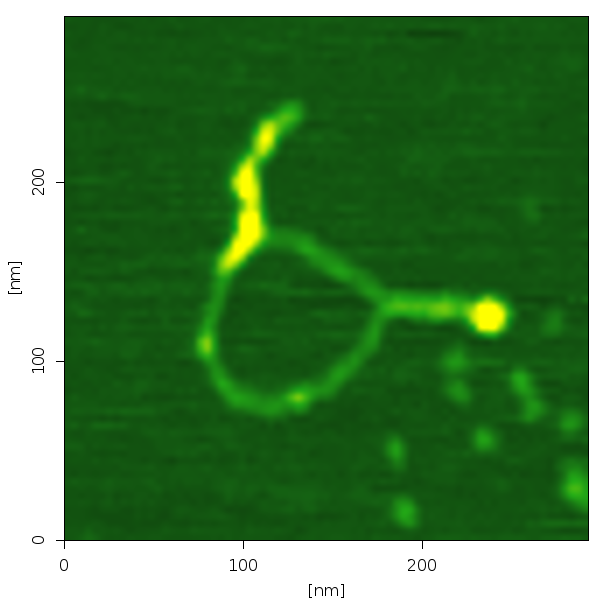

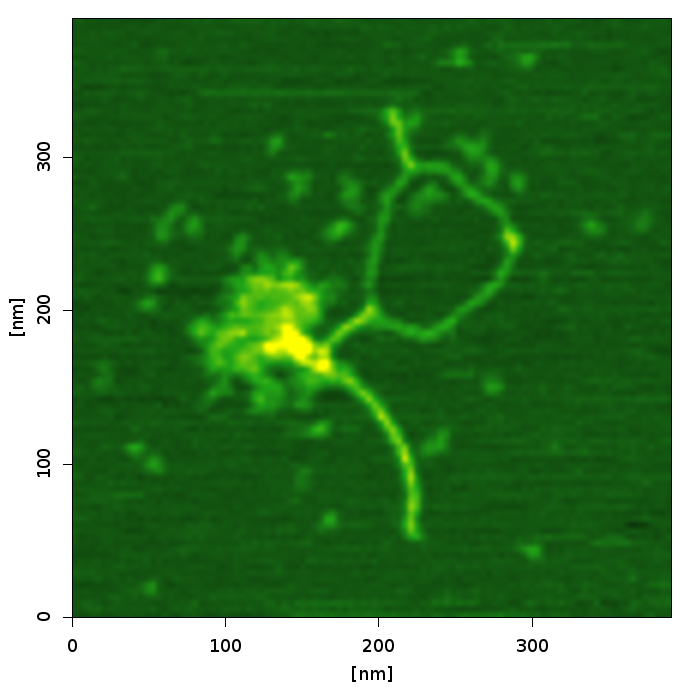

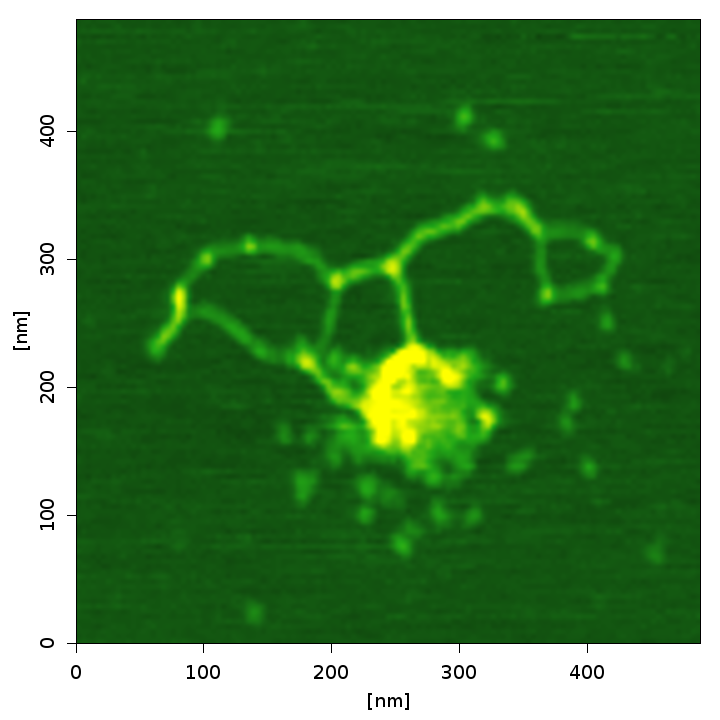

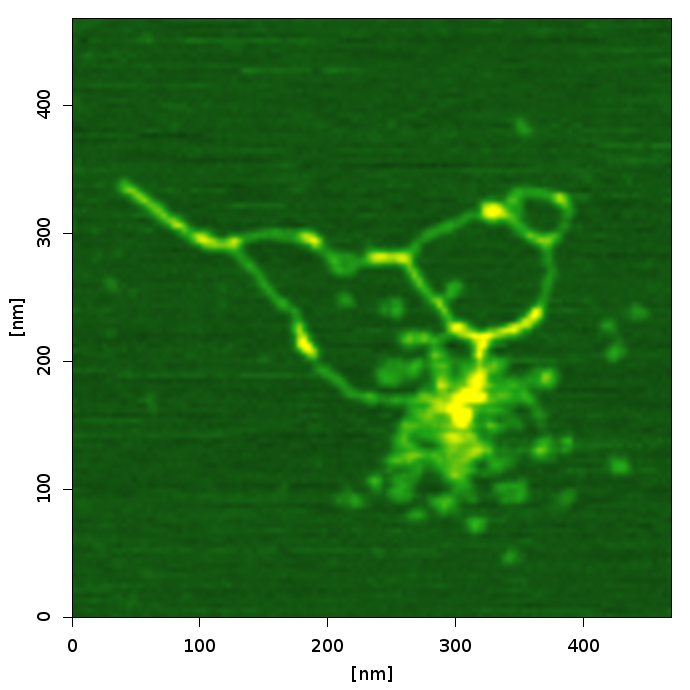

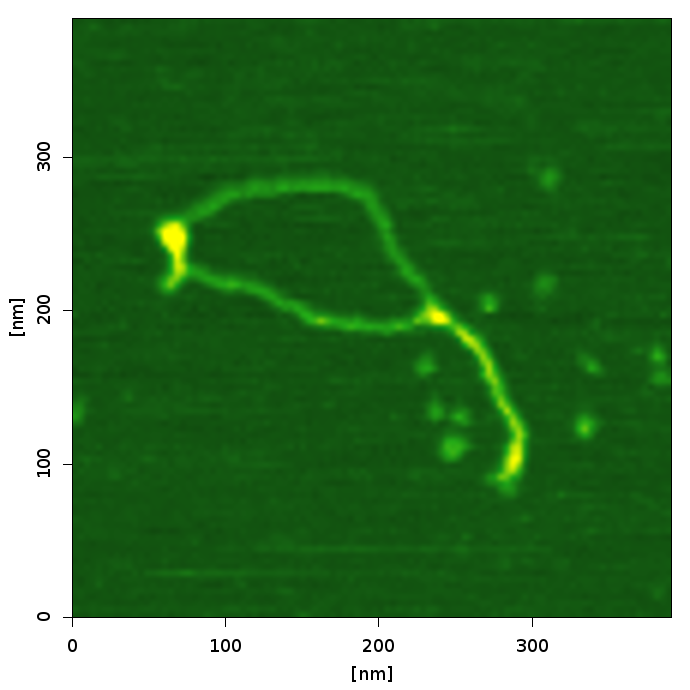

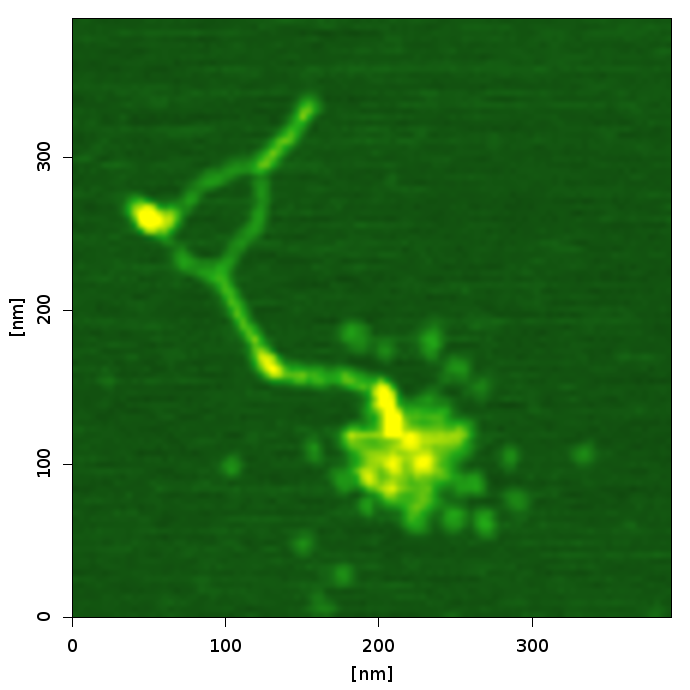

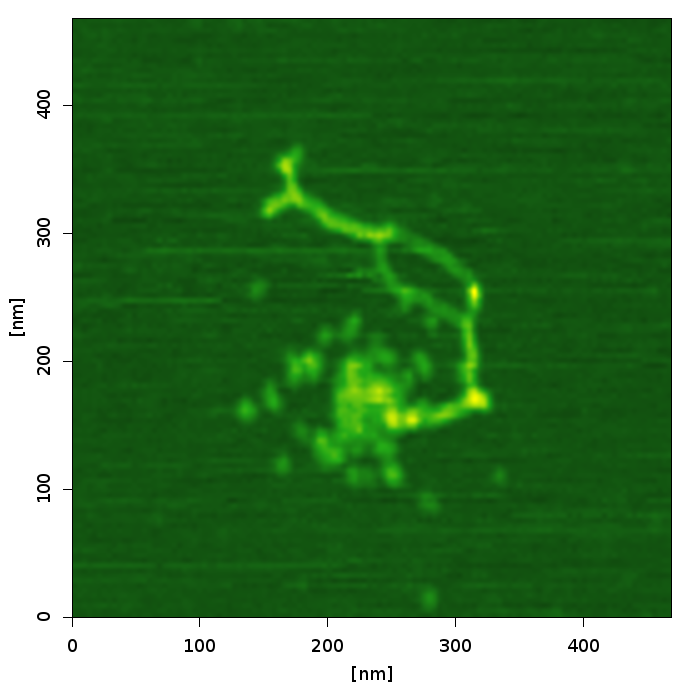

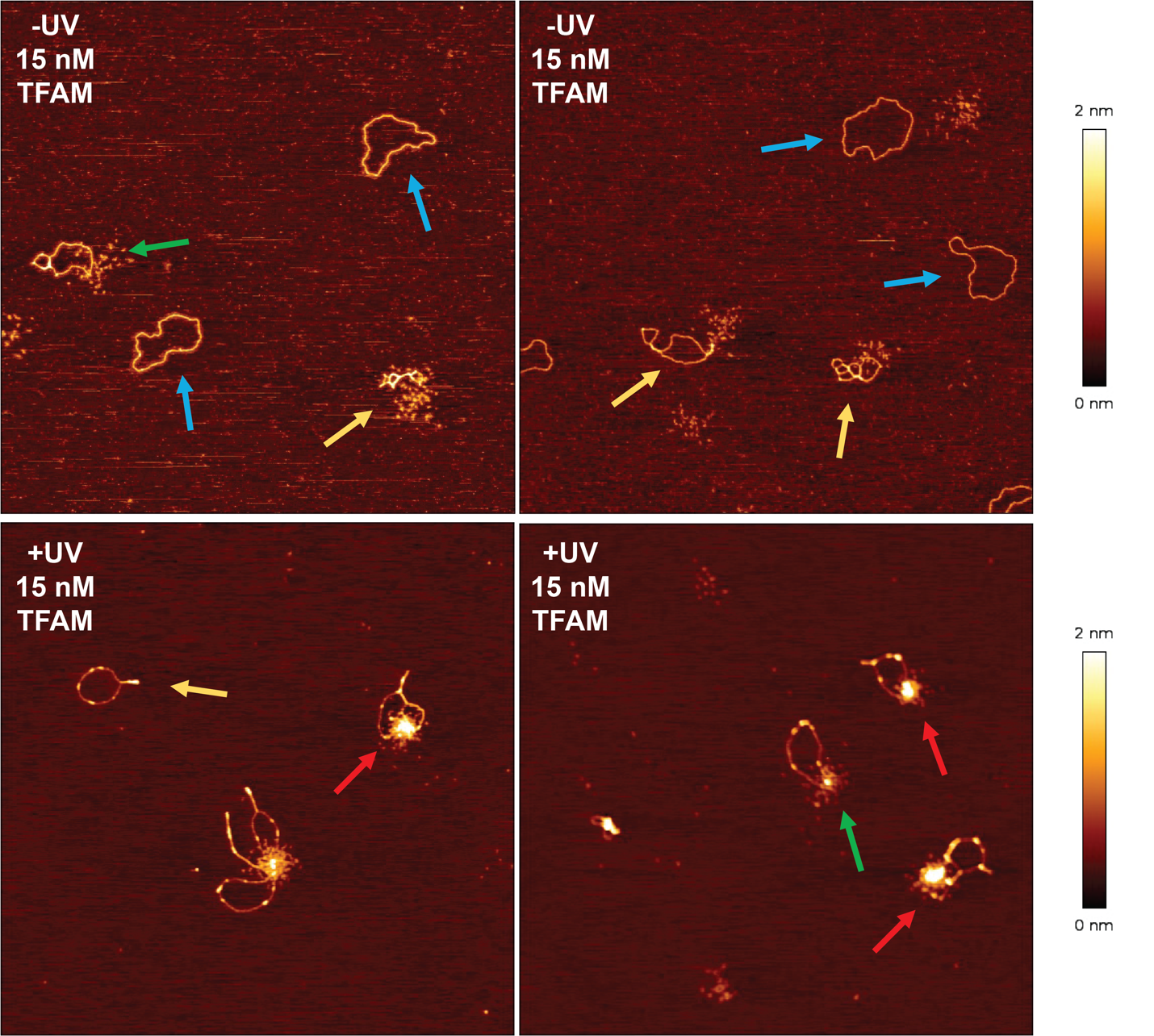

**Figure S18: AFM images of 15 nM TFAM incubated with either damaged or undamaged DNA.** The top panels show examples of 15 nM TFAM in the presence of undamaged (-UV) pUC19 DNA and the bottom panels show examples of 15 nM TFAM in the presence of damaged (+UV) pUC19 DNA. Colored arrows on the AFM images represent the different TFAM-DNA complexes categorizations dispersed (green), intermediate (yellow), punctate (red), and protein free DNA (blue). Each AFM image is 2x2 μm and 512x512 size pixels.

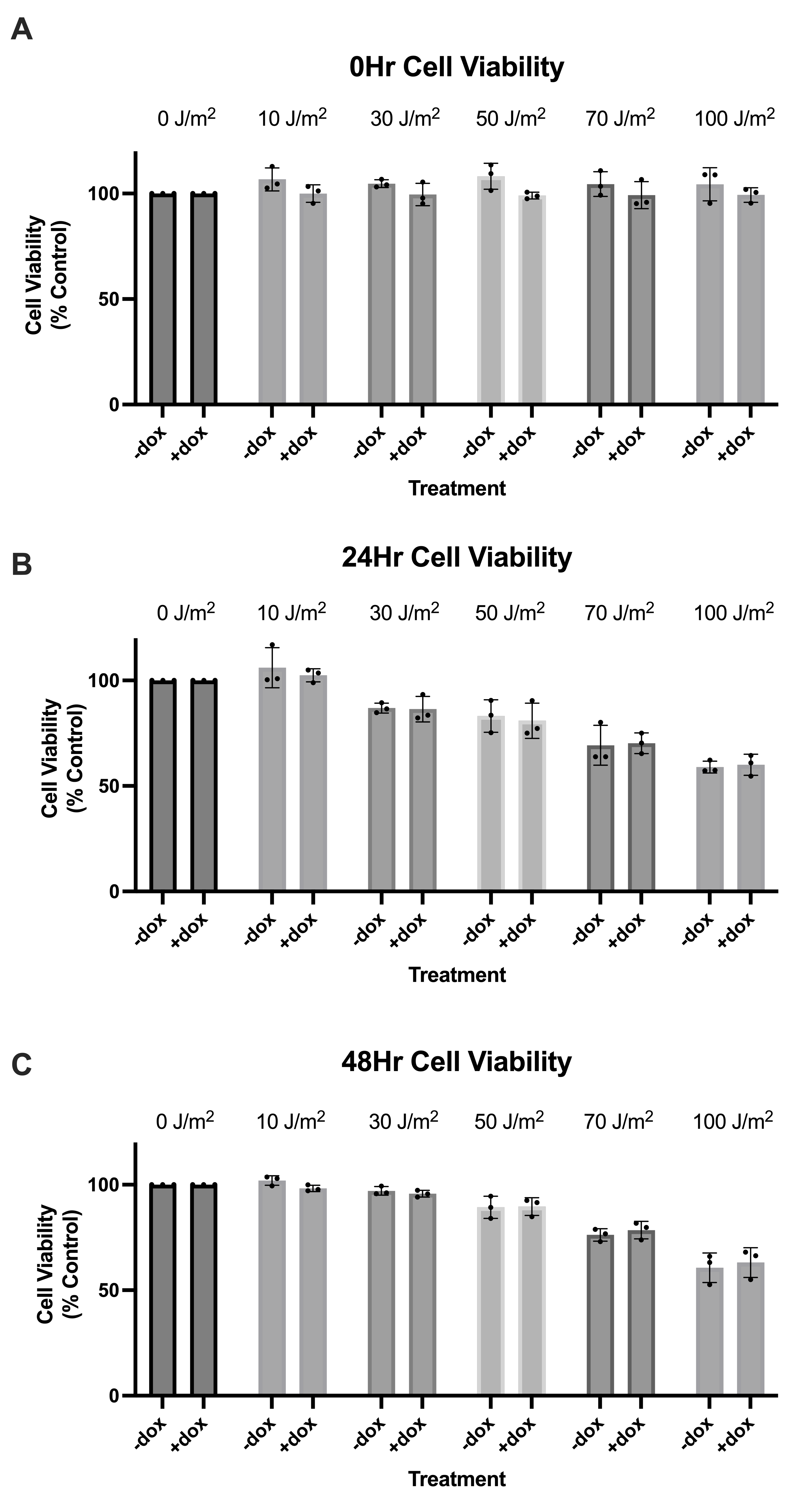

**Figure S19: TFAM overexpression does not protect cells from loss of cell viability from UVC exposures.** Cell viability following exposure to UVC normalized to the control for each replicate (n=3), determined using a resazurin cell viability assay. For all panels, x-axes represent the doxycycline treatment conditions across a range UVC dose delivered to the cells in J/m^2^. Y-axes represent the fluorescence readout normalized as percent control. Data was analyzed via two-way ANOVA. **A)** Cell viability immediately after exposure to UVC (UVC dose: p=0.8087, doxycycline treatment: 0.0029, interaction: p=0.6965). **B)** Cell viability 24 hours after exposure to UVC (UVC dose: p<0.0001, doxycycline treatment: 0.7183, interaction: p=0.9790). **C)** Cell viability 48 hours after exposure to UVC (UVC dose: p<0.0001, doxycycline treatment: 0.9968, interaction: p=0.7527).
